# Identification and quantification of human relaxin proteins by immunoaffinity-mass spectrometry

**DOI:** 10.1101/2024.01.12.575440

**Authors:** Yasmine Rais, Andrei P. Drabovich

## Abstract

The human relaxins belong to the Insulin/IGF/Relaxin superfamily of peptide hormones, and their physiological function is primarily associated with reproduction. In this study, we focused on a prostate tissue-specific relaxin RLN1 (REL1_HUMAN protein), and a broader tissue specificity RLN2 (REL2_HUMAN). Due to their structural similarity, REL1 and REL2 proteins were collectively named a ‘human relaxin protein’ in previous studies and were exclusively measured by immunoassays. We hypothesized that the highly selective and sensitive immunoaffinity-selected reaction monitoring (IA-SRM) assays could reveal the identity and concentration of REL1 and REL2 in biological samples and facilitate evaluation of these proteins for diagnostic applications. RT-PCR revealed the high levels of RLN1 and RLN2 transcripts in prostate and breast cancer cell lines. However, no endogenous prorelaxin-1 or mature REL1 were detected by IA-SRM in numerous biological samples. IA-SRM assay of REL2 revealed its undetectable levels (<9 pg/mL) in control female and male sera, relatively high levels of REL2 in maternal sera (median 331 pg/mL, 120 patients), and a biphasic expression of REL2 across the gestational weeks. IA-SRM assays discovered potential cross-reactivity and false-positive measurements for relaxin immunoassays. The developed IA-SRM assays will facilitate investigation of physiological and pathological roles of REL1 and REL2 peptide hormones.

## INTRODUCTION

The human relaxin genes belong to the Insulin/Insulin-like growth factor/Relaxin superfamily of ten peptide hormones that share high structural similarity despite relatively low sequence homology. These peptides include human insulin (INSL), insulin-like growth factors IGF1 and IGF2, relaxins RLN1, RLN2 and RLN3, and the insulin-like peptides INSL3, INSL4, INSL5, and INSL6^1^. Based on the Human Protein Atlas classification of the RNA-derived tissue specificity, several genes of the insulin/IGF/Relaxin superfamily exhibit the tissue-specific expression profiles. This includes the categories of the highly tissue-enriched RLN1 (prostate), INS (pancreas), IGF2 (placenta), INSL4 (placenta), INSL5 (interstine), a category of the group enriched RLN3 (parathyroid gland, testis), INSL3 (ovary, testis), and INSL6 (retina, testis), and a category of the tissue-enhanced RLN2 (fallopian tubes) and IGF1 (cervix). The high tissue specificity makes these genes and proteins promising biomarkers^2^.

We have previously evaluated a number of prostate tissue-specific proteins including KLK4, TGM4, and TMPRSS2-ERG fusion protein, but revealed their insufficient diagnostic performance^3-5^. Evaluation of the Human Protein Atlas for the top prostate-specific transcripts revealed REL1 (RLN1 gene) as the only secreted protein which has never been evaluated as a prostate cancer marker. The literature review revealed limited and contradictory information about REL1 measurements. Due to potential cross-reactivity of anti-REL1 and anti-REL2 antibodies and immunoassays, these two human genes and proteins were collectively referred to as a “human relaxin protein”.

Similar to insulin, prorelaxin proteins are cleaved into the mature REL1_HUMAN and REL2_HUMAN secreted peptides (∼6 kDa), as well as the connecting C-peptides^6-8^. The mature relaxins share 82% homology and comprise of A and B chains connected by two interchain disulfide bonds (**Figure 1A**). Relaxin protein sequence analysis (**Figure 1B**) reveals numerous lysine and arginine residues, short tryptic peptides, and peptides with numerous post-translational modifications. Low expression levels and lack of robust proteotypic peptides make REL1 and REL2 challenging proteins for identification and quantification with the conventional proteomic approaches and may explain inconsistent information about their expression. For example, a highly prostate-tissue specific RLN1 was listed by Uniprot^9^, NextProt^10^ and the Human Protein Atlas^11^ as a gene with evidence at protein level and as a plasma secreted protein, without experimental data (detection in blood plasma by immunoassays, mass spectrometry, or proximity extension assays) to support those observations. Likewise, Peptide Atlas^12^ presented identifications of the signal peptide and connecting C-peptide sequences in some studies, while the tryptic peptides of the mature REL1 have never been experimentally observed.

**Figure 1.**
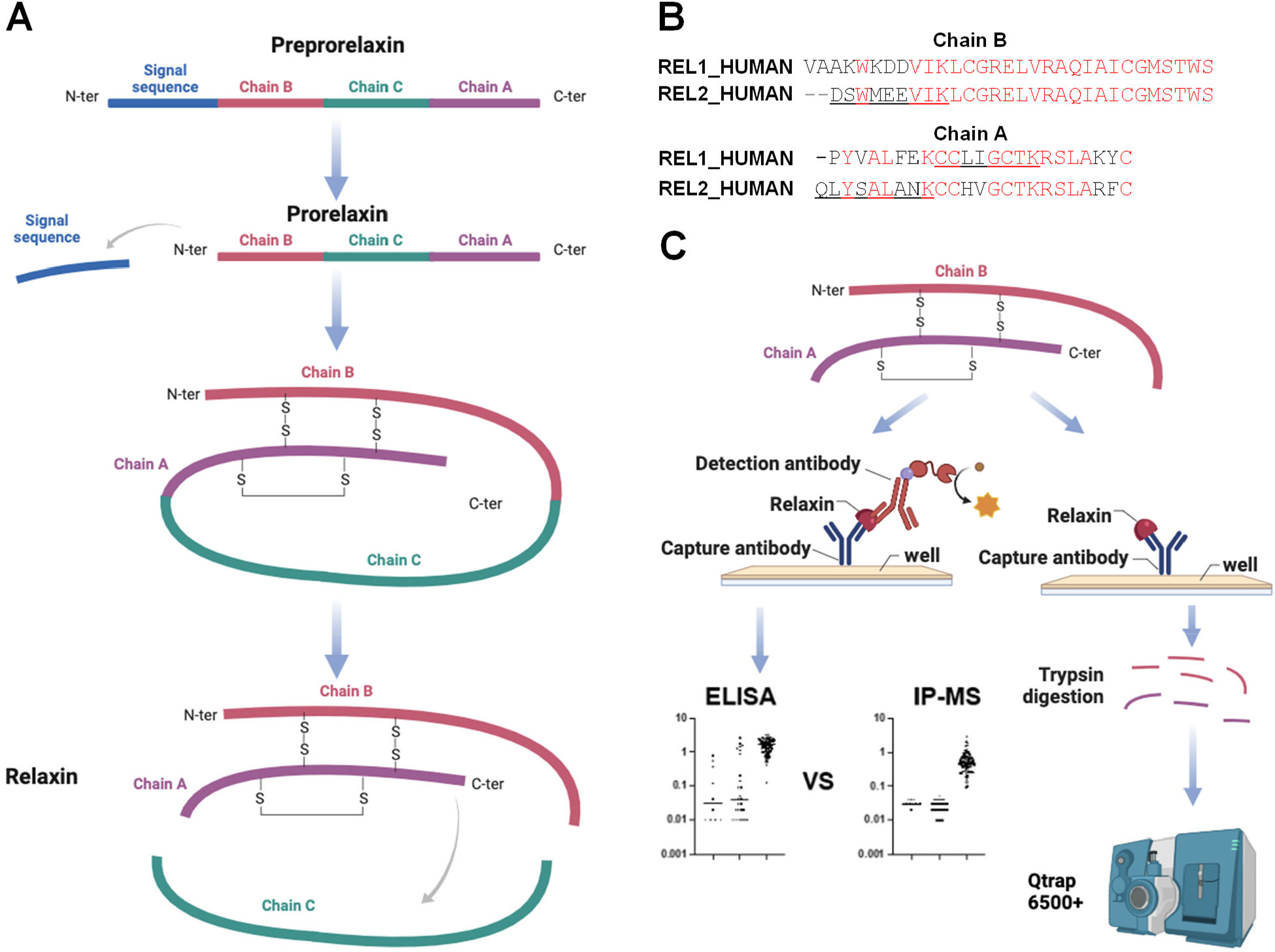
Experimental workflow for quantification of the human relaxin proteins. **(A)** Similar to insulin, preprorelaxin H2 (REL2_HUMAN; P04090-1) is translated as a 185 amino acid peptide whose signal peptide facilitates translocation into the endoplasmic reticulum (ER) and is proteolytically removed. ER-mediated oxidative folding forms disulfide bonds, and trans-Golgi processing removes the connecting C-peptide to generate a mature form of REL2 with chains B (29 aa) and A (24 aa). **(B)** Alignment of REL1 and REL2 chains B and A reveals their high sequence similarity. **(C)** Either recombinant or endogenous relaxin proteins were enriched from serum or biological samples using the identical capture antibody coated onto 96-well microplates. One of the microplates was utilized for ELISA immunoassay measurements, while another microplate was subjected to proteomic samples preparation, trypsin digestion of relaxin-antibody complexes to release prototypic peptides, and identification and quantification of tryptic peptides by nanoflow liquid chromatography – mass spectrometry.

In this study, we aimed at developing the highly selective and sensitive immunoaffinity-selected reaction monitoring (IA-SRM) assays as alternative tools to validate the human relaxin immunoassays (**Figure 1C**), reveal the identity and concentration of REL1 and REL2 in biological samples, and facilitate evaluation of these proteins for diagnostic applications.

## EXPERIMENTAL PROCEDURES

### Chemicals, reagents, and samples

Dithiothreitol, iodoacetamide, and trifluoracetic acid (TFA) were purchased from Thermo Fisher Scientific (Burlington, ON, Canada). Mass spectrometry-grade acetonitrile (ACN) and water were obtained from Fisher Scientific (Fair Lawn, NJ). Formic acid (FA), dimethyl sulfoxide (DMSO), and dimethylated SOLu-trypsin were obtained from Sigma-Aldrich (Oakville, ON). Stable SpikeTides_L peptides were obtained from JPT Peptide Technologies (Germany). The use of archived deidentified serum and seminal plasma samples was approved by the Health Research Ethics Board of Alberta (#HREBA.CC-22-0056) and the University of Alberta (#Pro00104098).

### Clinical samples

Leftover deidentified healthy female and male samples were obtained from the Alberta Health Services, and the use of these samples for our research study was approved by the Health Research Ethics Board of Alberta (HREBA.CC-22-0056). Leftover healthy normal maternal serum samples were previously collected for the study by Chan *et al* ^13^ and approved for research study by the University of Alberta’s Health Research Ethics Board. Briefly, maternal sera were selected from an archive of samples from women who elected to undergo second trimester (15–16 weeks) prenatal “triple screens” for trisomy 18, Down’s Syndrome, and open spina bifida. All women were ≥18 years of age and delivered at ≥22 weeks of gestation to live singletons without malformations ^13^. The use of these archived samples for our study has been approved by the Health Research Ethics Board of Alberta (#HREBA.CC-22-0056). Seminal plasma samples were obtained from the APCaRI Biorepository (https://apcari.ca) and their use was approved by the Health Research Ethics Board of Alberta (#HREBA.CC-14-0085 and HREBA.CC-22-0056).

### Cell culture

Cell lines were obtained from ATCC (https://www.atcc.org). Cell lines were cultured in a humidified incubator at 37 °C and 5% CO2 in T25 flasks to ∼80% confluency. A prostate cancer LNCaP cell line was cultured in RPMI 1640 medium (Gibco) media supplemented with 10% fetal bovine serum (Invitrogen) and 1% penicillin-streptomycin (Invitrogen). The breast cancer cell lines MDA-MB-231 and MCF-7 were cultured in Dulbecco’s modified Eagle’s medium (HyClone DMEM) supplemented with 10% v/v fetal bovine serum, 5% non-essential amino acids, 50 units/ml penicillin, and 50 μg/ml streptomycin. All cells were washed three times with phosphate-buffered saline (PBS), and snap-frozen at -20 °C until further analysis.

### Cell lysis

Cell pellets were exposed to 20 µl of EDTA-free protease inhibitor cocktail (Roche) to prevent protein degradation, lysed with 50 µl of 0.1% RapiGest SF (Waters), vortexed and probe sonicated at 20 kHz for 1 minute. Lysates were centrifugated at 16,000g and 4 °C for 10 min to remove cell debris. Total protein content in the lysates was determined using the Pierce BCA protein assay kit (Fisher Scientific).

### Cell line secretome

LNCaP cells growing in a serum-free medium were serum-deprived for 48 h. Conditioned medium (supernatant) with cell line secretome was collected and stored at -20 °C. Supernatant (5 ml) was added to the 0.5-1 kDa molecular cut-off membrane (Spectrum Laboratories) and dialyzed in 1 mM ammonium bicarbonate with two buffer exchanges for 24 hours at 4° C. Dialyzed supernatant was concentrated 10-fold to 0.5 mL using a Speed Vacuum Concentrator.

### Centrifugal ultrafiltration of sera

Pooled maternal sera (500 μL) was diluted 10-fold with 10% ACN. The mixture of serum and ACN was placed on ice for 1 hour and then centrifuged at 5,000 rpm for 5 minutes. The supernatant was fractionated using a prepared low-protein binding Vivaspin-2 20K MWCO membrane filter at 4,000 g until 95% of the input serum had passed through the 20K (cellulose triacetate) filter. The concentrate and filtrate were subjected to immunoaffinity enrichment and Q-Exactive analysis.

### ELISA

Human REL1 DuoSet ELISA (DY3257) and Human REL2 DuoSet ELISA (DY2804-05) assays were purchased from R&D systems. The quantification of recombinant REL1 and recombinant REL2 was performed following the manufacturer’s protocol.

### SDS-PAGE

SDS-PAGE was performed using a BioRad Mini-Cell electrophoresis unit utilizing Mini-PROTEAN TGX Precast Gels (4–15% gradient acrylamide). The PageRule Protein (250 to 5 KDa) was used as a ladder. Under non-reducing conditions, Tris/Glycine/SDS buffer (#1610732EDU) was used as the buffer system at a constant voltage of 120 V for 80 min.

### RT-PCR

Total RNA was extracted from LNCaP, Hela and MCF-7 cells using TRIzoL (Thermo Fisher Scientific). RNA was reversely transcribed to cDNA via iScriptTM Reverse Transcription Supermix (Bio-Rad Laboratories). After quantification by NanoVue Plus spectrophotometer (GE Healthcare), 500 ng cDNA was utilized as a template for amplification of RLN1 and RLN2 using Hot Start Taq 2X Master Mix (New England Biolabs) and GeneAmp PCR System 2700 thermal cycler (Applied Biosystems). The forward and reverse PCR primers for RLN1 gene included 5’-CCTGTTCTTGTTCCACCTGC-3’ and 5’-AGCTCTGGTAATGATGGTTGC-3’ (Integrated DNA Technologies) and RLN2 primers were 5’-CGGACTCATGGATGGAGGAA -3’ (forward) and 5’-GTAGCTGTGGTAATGCTGGCC -3’ (reverse). The final volume was 25 μl, and an initial denaturation step of 95 °C for 5 min was followed by 35 cycles of 95 °C for 30 seconds 53 °C for 30 s and 72 °C for 30 seconds, and one cycle at 72 °C for 5 min. RLN1 and RLN2 cDNA were detected by 2% agarose gel electrophoresis.

### Immunoaffinity enrichment and proteomic sample preparation

Anti-REL1 rat monoclonal antibody (part # 843350), and anti-REL2 mouse monoclonal antibody (part # 844036) from the ELISA kits (R&D system) were used as capture antibodies for the enrichment of REL1 and REL2 proteins from cell lysate and clinical samples. 500 ng of each antibody was coated onto high-binding 96-well polystyrene microplates (#07000128; Greiner Bio-One) at 100 μl per well and incubated overnight at room temperature (RT). Following three washes with 0.1% Tween 20 in PBS (washing buffer; 200 μl each), the plate was blocked for 1 hour at RT with 200 μl of blocking buffer (1% BSA in wash buffer). After repeated washing, 100 μg of total cell lysate protein per well or 50 μl of clinical samples (blood serum and seminal plasma) were diluted with 50 μl of the dilution buffer (0.1% BSA in wash buffer, 0.2 μm-filtered) and added to microplates. Following a 2-hour incubation with shaking, the plate was washed three times with 200 μl of washing buffer and three times with 50 mM ammonium bicarbonate. The enriched proteins underwent reduction with dithiothreitol, alkylation with iodoacetamide, and digestion on the same plate using trypsin (0.5 μg/well). SpikeTides_L (**supplementary Table S1**) peptide standards (200 fmol/well) were added after digestion. Tryptic peptides were desalted and concentrated with C18 microextraction tips (Agilent).

### Development of SRM assays

Peptide Atlas^12^, neXtProt^10^ and our shotgun MS data were used to select proteotypic peptides for REL1 and REL2 proteins, including chain A and B tryptic peptides, as well as connecting chain C peptides representing immature relaxin forms. In total, 17 synthetic heavy isotope-labeled SpikeTides_L peptides were used as internal standards for the SRM assay development (**supplementary Table S1**). Charge states and collision energies were determined based on the SRM peak area, and peptides with poor performance were excluded. Light and heavy transitions of low-intensity and with interferences were eliminated upon analysis of recombinant relaxin. As a result, six unique REL1 and REL2 peptides with the three most intense and reproducible transitions per peptide light or heavy form were scheduled within a 2-minute acquisition window (**supplementary Tables S2, S3**). A scan time of 10 ms provided at least 20 points per peak. The correct identification of peptides was confirmed by examining the superposition of light and heavy transition peaks, peak shapes, and the order of y-ion transition intensities. The amounts of the endogenous light peptides were calculated using the peak area ratio of the spiked-in heavy peptide internal standards.

### Chromatography and targeted mass spectrometry

A quadrupole ion-trap mass spectrometer (SCIEX QTRAP 6500+) coupled to optiFlow Turbo V nanospray ion source and Eksigent Ekspert nanoLC 425 (SCIEX) was used for SRM assays. The tryptic peptides were loaded at 5 μl/min onto a C18 nano trap RP-1 column (Phenomenex, 10×0.075 mm, 5 μm) and separated with BioZen columns (Phenomenex Peptide Polar-C18, 150×0.075 mm, 2.6 μm, 100 Å) and 40 min gradients (300 nL/min). The gradient started with 5% buffer B and ramped to 50% buffer B over 11 min, then to 90% buffer B within 2 min, and continued for 2 min, then to 5% buffer B in 1 min and continued for 24 min. Nanospray ion source and QTRAP 6500+ parameters included 3,100 V nanospray, 150°C heater temperature, 9 psi for gas 1 (N2), 0 psi gas 2; 30 psi curtain gas (N2), and 80 V declustering potential. Q-Exactive Hybrid Quadrupole-Orbitrap (Thermo Scientific) coupled to Easy-Spray source and EASY-nLC 1000 (Thermo Scientific) was used for analysis of seminal plasma samples. The mobile phase consisted of 0.1% formic acid in water (buffer A) and 0.1% formic acid in acetonitrile (buffer B). Acclaim PepMap 100 nanoViper C18 (Thermo Scientific, 100 μm×2 cm, 5 μm, 100 Å) was used as a trap column, while EASY-Spray C18 (Thermo Scientific, 15 cm×75 μm, 3 μm) was used as an analytical column. An 18-min gradient (400 nL/min) started with 0% buffer B and ramped to 50% buffer B over 15 min, then to 100% buffer B within 1 min, and continued for 2 min. PRM scans were performed at 17.5 K resolution with 27% normalized collision energy. Automatic Gain Control target value was set to 3×10^6^ (100 ms injection time; 2.0 m/z isolation width). The performance of the nanoLC-MS systems was assessed every six runs with BSA digest solution (10 μl of 10 fmol/μl).

### Shotgun mass spectrometry

REL1 proteotypic peptides were identified with nanoflow liquid chromatography-tandem mass spectrometry (LC-MS/MS) and an Orbitrap Elite mass spectrometer equipped with Easy-Spray source and EASY-nLC II system (Thermo Scientific). Peptides were separated at 300 nL/min with a 2-hour gradient: 95% buffer A (0.1% formic acid in water) and 5% buffer B (0.1% formic acid in ACN) for 5 min, 5-35% B for 95 min, 35-65% B for 10 min, 65-100% B for 1 min and 100% B for 9 min. The analysis included profile MS1 scans (400−1250 m/z; 60 K resolution), followed by top 20 ion trap MS/MS scans, acquired at 33% normalized collision energy. FTMS ion count was set to 1×10^6^ with an injection time of 200 ms, while MS/MS scans were set to 9,000 counts and 100 ms injection time. MS/MS acquisition settings included 500 minimum signal threshold, 2 m/z isolation width, 10 ms activation time, and 60 s dynamic exclusion. Monoisotopic precursor selection was enabled, and +1 and unknown charge states were rejected. Instrument parameters included 230°C capillary temperature and 2.0 kV spray voltage. Raw files were searched using MaxQuant v1.6.3.4 against the reviewed Uniprot database. Modifications considered constant cysteine carbamidomethylation and variable methionine oxidation, N-terminal acetylation, N and Q deamidation, N-term pyro-cmC, and pyro-Glu.

## RESULTS

### Development of SRM assays for quantification of REL1 and REL2 proteins

Due to their short sequence and numerous lysine and arginine residues, REL1_HUMAN and REL2_HUMAN proteins have only a few trypsin-cleavable proteotypic peptides. To select the best proteotypic peptides, we digested by trypsin the recombinant REL1 and REL2 and identified their peptides by shotgun mass spectrometry (nanoLC-nanoESI-MS/MS). Likewise, we obtained several heavy isotope-labeled synthetic peptides and analyzed them by targeted mass spectrometry (nanoLC-nanoESI-SRM). As a result, we selected proteotypic peptides for quantification of REL1 (CCLIGCTK) and REL2 (QLYSALANK and DSWMEEVIK) (**Figure 2**). We also evaluated post-translational modifications of the N-terminal cysteine (S-carbamoylmethylcysteine cyclization, or pyro-cmc, denoted as “c”) and glutamine (pyroglutamic acid, or pyro-Glu, denoted as “q”). We detected consistent formation of N-term pyro-cmc in cCLIGCTK (71.4% of total amount) and N-term pyro-Glu in qLYSALANK (21.5% of total amount) in synthetic peptides.

**Figure 2.**
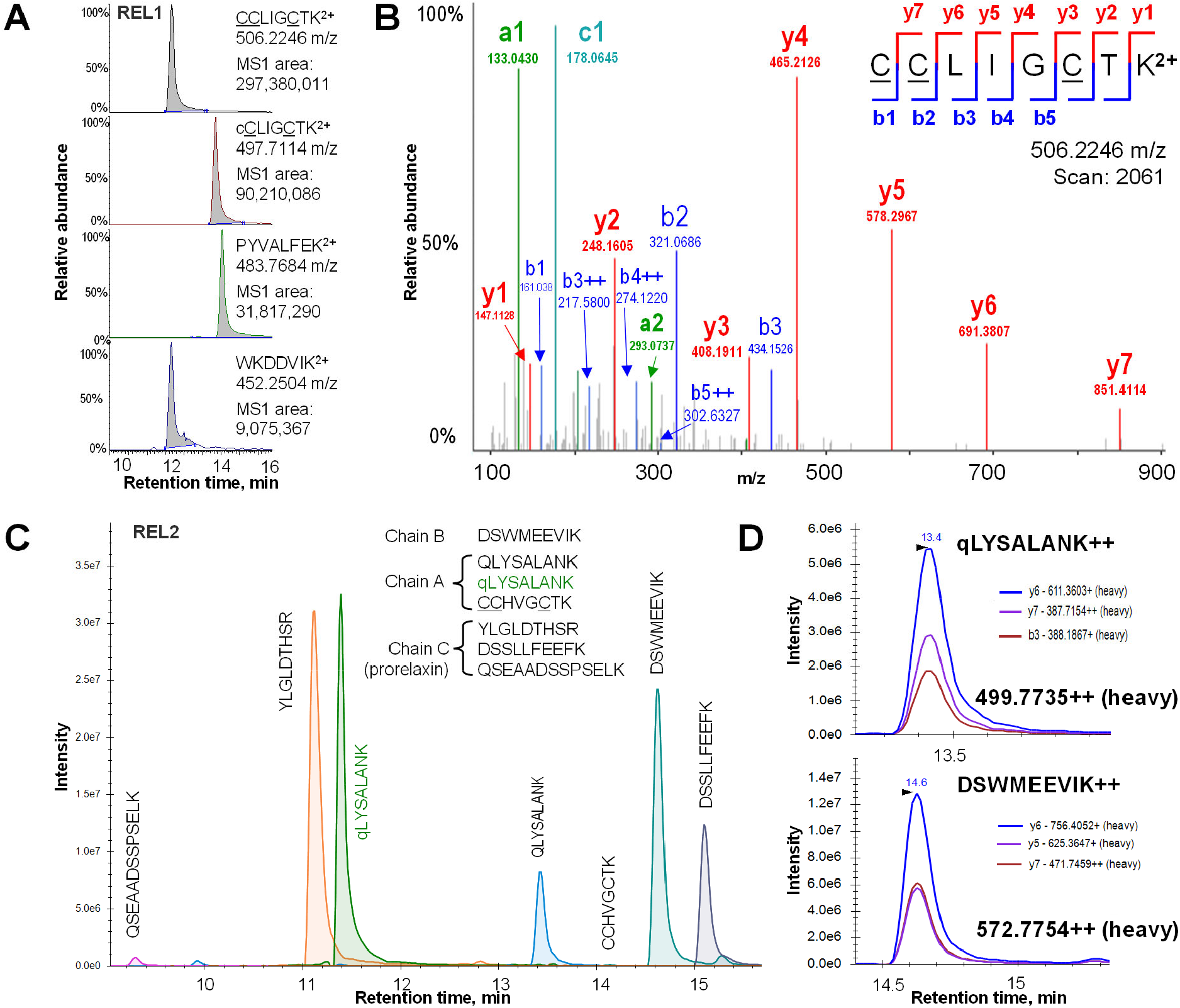
Selection of proteotypic peptides for REL1 and REL2 proteins. **(A)** The recombinant mature REL1 was digested by trypsin, and peptides were analyzed by nanoLC-shotgun MS/MS. <u>CC</u>LIG<u>C</u>TK peptide was selected as the most intense MS1 peptide, while MS/MS search provided fragments with the highest signal-to-noise ratio (y4, y5, and y6). **(B)** Tryptic peptides of mature REL2 and a C-peptide were selected, synthesized as heavy isotope-labeled peptides, and analyzed by nanoLC-targeted MS. DSWMEEVK and qLYSALANK peptides demonstrated the highest intensity and were selected as proteotypic peptides to quantify mature REL2. RT, retention time; <u>C</u>, S-carbamidomethylated cysteine; c, N-term pyro-carbamidomethylated cysteine, or pyro-cmC; q, N-term pyroglutamic acid, or pyro-Glu.

### Evaluation of immunoassays and development of IA-SRM assays

Colorimetric ELISA immunoassay revealed LODs of 69.6 ng/mL and 4.1 pg/ml for the recombinant mature REL1 and REL2, respectively (**Figure 3**). No substantial cross-reactivities of REL1 with an REL2 immunoassay and REL2 with an REL1 immunoassay were observed (**supplementary Fig. S1)**.

**Figure 3.**
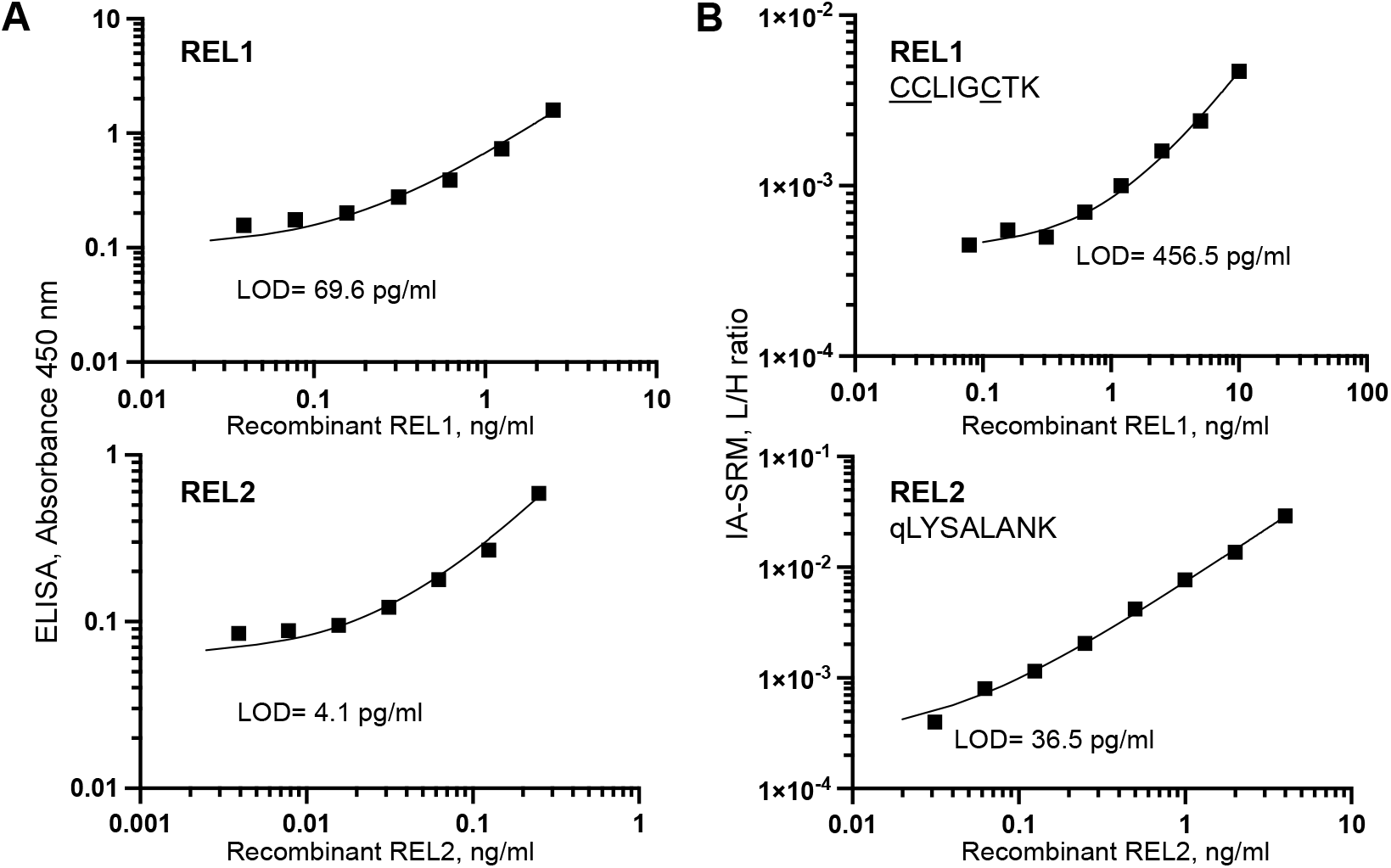
Evaluation of colorimetric ELISA and IA-SRM assays to quantify the recombinant mature REL1 and REL2 proteins. Each dilution was analyzed in triplicates by ELISA. Each analytical (full process) sample was analyzed in technical duplicates by IA-SRM. LODs were defined as the lowest concentration measured with CV<20%. Recombinant REL1 measured with ELISA and IA-SRM assays revealed LODs of 69.6 pg/mL and 456.5 pg/mL, respectively. Recombinant REL2 measured with ELISA and IA-SRM (qLYSALANK; 3 transitions) revealed LODs of 4.1 pg/mL and 36.5 pg/mL, respectively.

To develop IA-SRM assays, we used REL1 and REL2 capture antibodies provided with the ELISA immunoassay kits. Briefly, antibodies were coated onto microplates, recombinant REL1 and REL2 proteins were enriched, digested by trypsin, and the proteotypic light endogenous peptides and heavy internal standard peptides were quantified by SRM. IA-SRM assays revealed LODs of 457 pg/mL for REL1 (<u>CC</u>LIG<u>C</u>TK peptide) and 36.5 pg/mL (qLYSALANK peptide) for REL2 (**Figure 3**). The difference in immunoassay and IA-SRM assay sensitivity could be attributed to the higher affinity of the anti-REL2 antibody and better ionization of REL2 proteotypic peptides. As an additional validation step, we confirmed that denatured recombinant REL1 could not be detected by IA-SRM or ELISA (**supplementary Fig. S2**).

### Measurement of REL1 and REL2 in cell lines

The RNA sequencing data of the Cancer Cell Line Encyclopedia (the Dependency Map portal)^14^ revealed the highest levels of RLN1 and RLN2 expression in an LNCaP prostate cancer cell line. Using RT-PCR, we experimentally confirmed the expression of RLN1 and RLN2 transcripts (**Figure 4**). The ELISA measurements revealed REL2 in LNCaP cell lysate and cell media (9.3 pg/ml and 9.1 pg/mL, respectively), but no REL1 expression (**Table 1**). To facilitate detection of REL1, LNCaP cells were grown in serum-free media, and the medium was collected, dialyzed, and partially lyophilized to decrease the volume 10-fold and concentrate proteins. However, no endogenous REL1 was detected by ELISA. Even though ‘a human relaxin’ protein has previously been associated with breast cancer^15,16^, we have also demonstrated that REL1 was not detectable in either MDA-MB-231 or MCF-7 breast cancer cell lines (**supplementary Table S4**).

**Table 1.** Quantification of REL1 and REL2 proteins by ELISA in LNCaP cell lysate and cell culture media. LODs of REL1 and REL2 ELISA were 69.5 and 4.1 pg/ml, respectively. Secreted prostate-specific antigen (mature KLK3_HUMAN protein) was used as a positive control for sample treatment protocols. The samples were analyzed in duplicates.

| Sample | REL1 | REL2, pg/mL | KLK3, ng/mL |
| --- | --- | --- | --- |
| LNCaP cell lysate | < LOD | 9.1 | 43 |
| LNCaP cell culture medium (before dialysis) | < LOD | < LOD | 0.61 |
| LNCaP cell culture medium (after dialysis) | < LOD | 9.3 | 1.6 |

**Figure 4.**
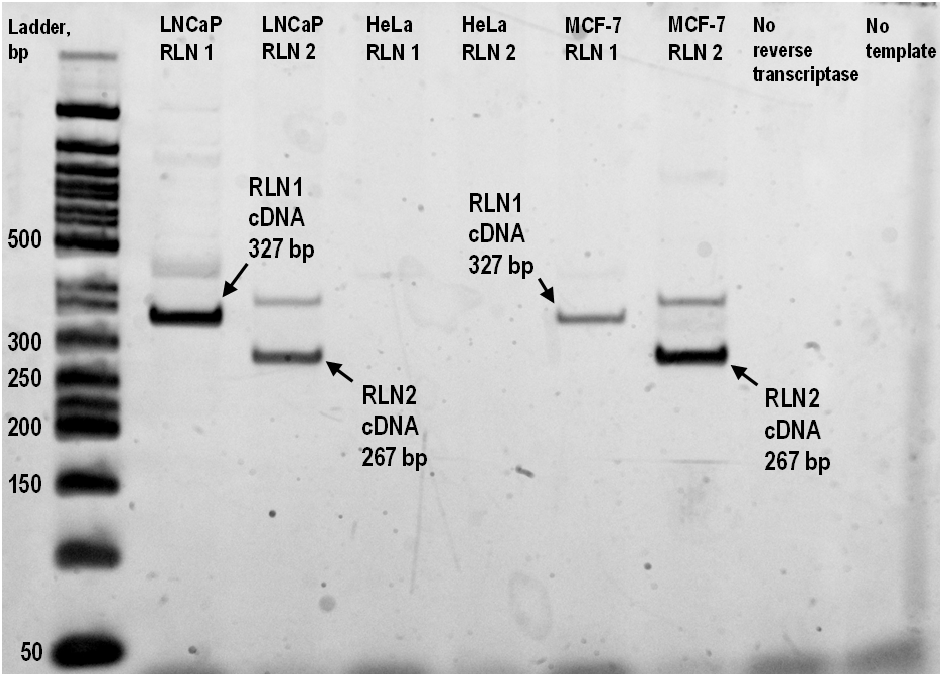
Expression of RLN1 and RLN2 mRNA transcripts in cell lines. RT-PCR revealed expression of RLN1 and RLN2 mRNA in a prostate-cancer cell line LNCaP, as confirmed by the agarose gel electrophoresis detection of cDNA products of predicted length. Cervical cancer HeLa and breast cancer MCF-7 cell lines were used as negative and positive controls, respectively, as suggested by the Cancer Cell Line Encyclopedia data.

### Measurement of REL1 and REL2 in seminal plasma

Previous studies demonstrated that the “human relaxin protein” (potentially a mixture of REL1 and REL2) was detected in prostate tissues and seminal plasma^17-22^. Here, we measured REL1 and REL2 by ELISA and IA-SRM in 18 seminal plasma samples and detected REL2 but not REL1 (**Figure 5**). Seminal plasma levels of REL2 (as high as ∼1 ng/ml) agreed with the previous studies (1.6 ng/mL)^20^. IA-SRM assay independently confirmed the expression of the endogenous REL2 in several samples. Our results suggested that the endogenous ‘human relaxin’ in seminal plasma was exclusively REL2, but not REL1. Further studies with ultrasensitive immunoassays^23^ could confirm potential REL1 expression at the levels below 70 pg/mL.

**Figure 5.**
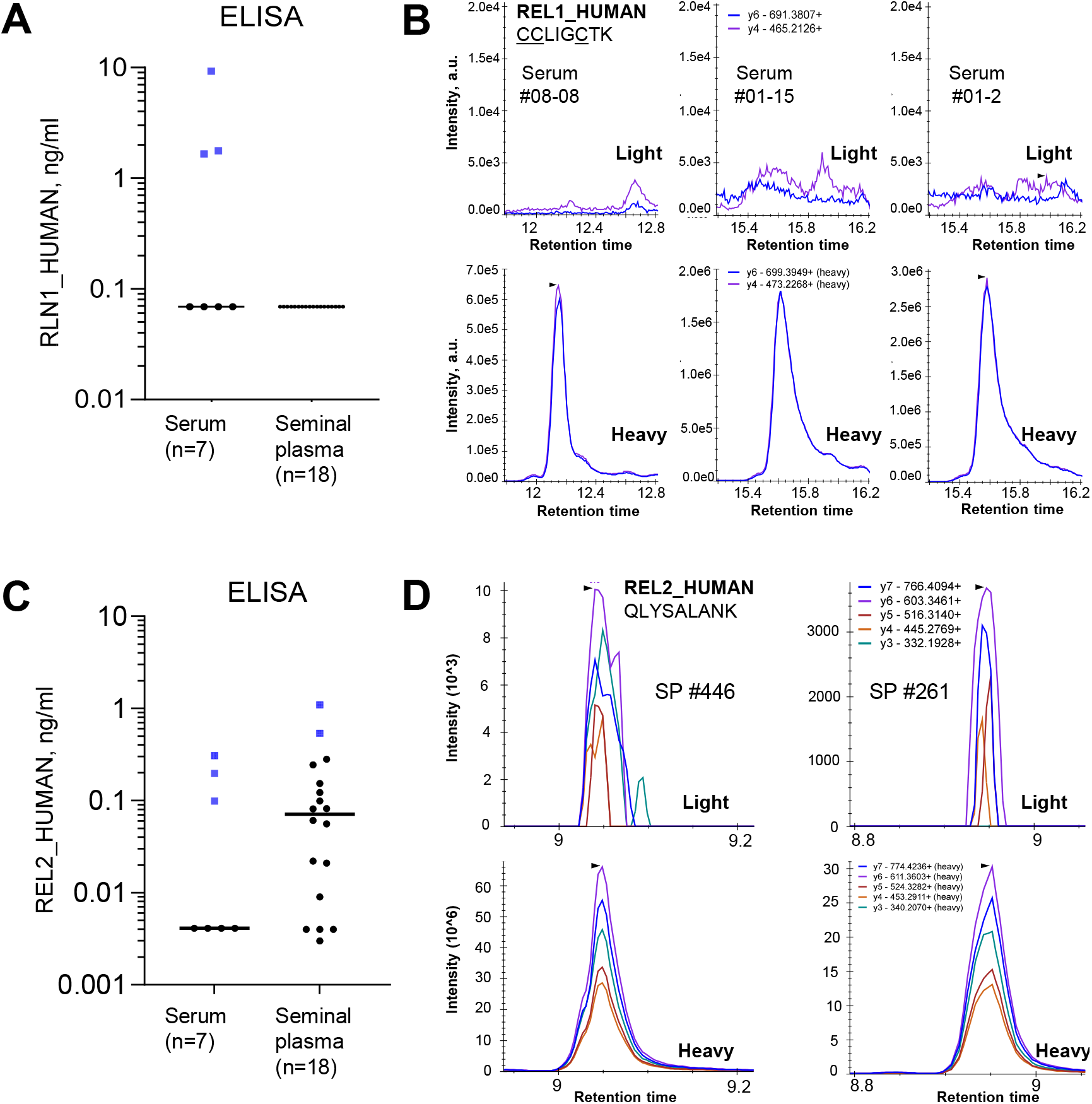
Measurement of endogenous REL1 and REL2 proteins in serum and seminal plasma samples with ELISA and targeted IA-SRM assays. Serum and seminal plasma samples positive on ELISA (blue squares) were subjected to IA-SRM measurements. **(A)** High levels of REL1 were detected with ELISA in three serum samples #08-08, #01-15, and #01-2. No REL1 protein was detected in seminal plasma. **(B)** IA-SRM measurements revealed background noise, but not any endogenous REL1 in three serum samples positive by ELISA. **(C)** Positive signal for REL2 protein was detected in serum samples #08-08, #01-15, and #01-2 by ELISA, but not IA-SRM. **(D)** REL2 ELISA-positive seminal plasma samples #446 and #261 were positive by IA-SRM.

### Measurement of REL1 and REL2 in blood serum

The “human relaxin protein” has previously been detected by ELISA in serum of healthy individuals and patients with prostate cancer and heart failure^24-29^. Here, we detected by ELISA the relatively high levels of REL1 (as high as ∼7,000 pg/ml) and REL2 (as high as ∼300 pg/ml) in several serum samples of healthy individuals (**Figure 5A**,**C**). IA-SRM measurements confirmed only REL2 in seminal plasma (**Figure 5D**). These results indicated potential cross-reactivity and false-positive identifications for REL1 and REL2 immunoassays in serum samples.

### Evaluation of REL1 and REL2 in maternal sera

Relatively high levels of relaxin (2.2 ng/ml) have previously been detected in maternal sera^30^. We hypothesized that maternal sera samples with high REL2 levels could also have measurable amounts of REL1. As a result, ELISA measurements revealed high levels of REL1 and REL2 in some maternal sera samples, while only REL2 was detected by IA-SRM (**supplementary Fig. S3**). In addition, we attempted to identify endogenous REL1 and REL2 (including their potential unexpected modifications) by shotgun LC-MS/MS. Maternal sera samples expressing high levels of REL2 were subjected to protein precipitation by ethanol and centrifugal filter purification (0.5-1 kDa cut-off) to remove high-abundance serum proteins (**supplementary Figs. S4** and **S5**). Following IA enrichment and trypsin digestion, an MS1 peak of the N-term pyroGlu-modified peptide qLYSALANK of REL2 was detected in maternal sera samples at 5 ppm mass tolerance but was not selected for MS/MS fragmentation. As a positive control, we analyzed recombinant REL2 (2 ng/mL) by IA-MS/MS using Thermo Q Exactive and a more sensitive Orbitrap Fusion Lumos Tribrid instrument. Both instruments detected MS1 peak of qLYSALANK, but neither instrument selected this peptide for MS/MS fragmentation. We concluded that the endogenous levels of REL2 are too low to be identifiable by the conventional shotgun proteomic approaches even after IA enrichment. These experiments justified the use of targeted MS to detect the endogenous REL2 following IA enrichment.

### Measurement of REL2 in maternal sera collected at different gestational weeks

Subsequently, we measured by ELISA and IA-SRM REL2 protein in a large set of maternal sera samples collected at different gestational weeks (N=120, weeks 13-29), as well as control female (N=30) and male (N=10) sera (**supplementary Table S5**). In this experiment, the IA-SRM assay LOD (S/N>3) has been further improved to 9.4 pg/mL (relative to 36.5 pg/mL reported in **Figure 2**) through the use of a higher amount of capture antibody (500 versus 300 ng/well), two additional calibration points (15.6 and 7.8 pg/mL), two quantifiable transitions with the lowest background (y6^1+^ and y7^2+^), and a sigmoidal curve fitting of a calibration curve (**supplementary Fig. S6** and **supplementary Table S6**). IA-SRM revealed a median concentration of 331 pg/mL for REL2, with concentrations ranging from 75 to 1,300 pg/mL (**Figure 6** and **supplementary Tables S7, S8**). The endogenous REL2 protein was detected only in a few female and male sera samples and at the levels approaching IA-SRM LOD (9.4 pg/mL). REL2 ELISA measurements provided median 1.6 ng/mL and revealed a reasonable correlation (r^2^=0.94) with qLYSALANK peptide IA-SRM levels (**supplementary Fig. S7** and **supplementary Table S9**). In addition, ELISA detected high concentrations of REL2 in some female and male samples (**Figure 6C** and **supplementary Table S10**). Evaluation of REL2 across gestational weeks revealed potential biphasic trend with a local minimum at 20 weeks (**Figure 6B, D** and **supplementary Table S11**).

**Figure 6.**
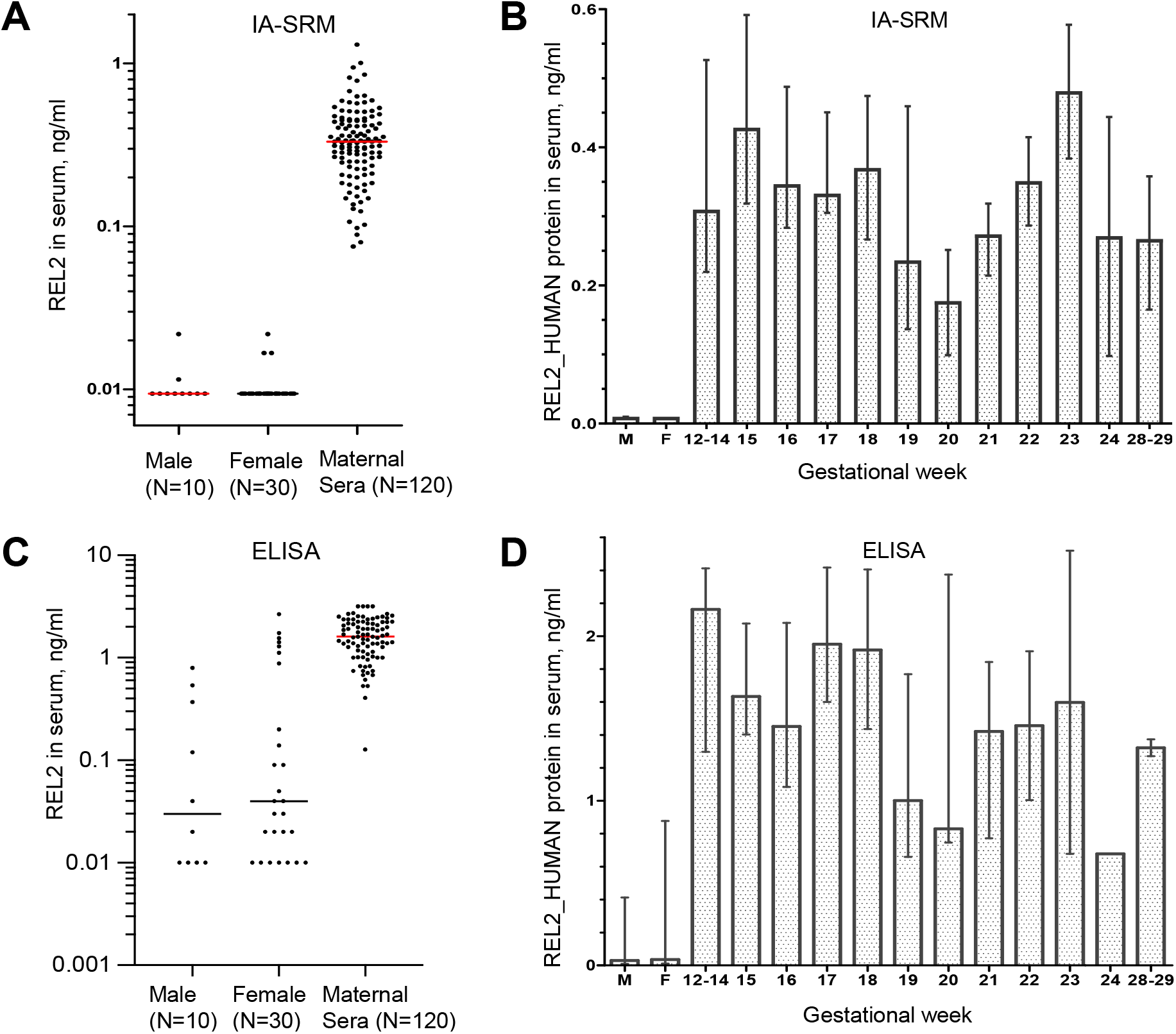
Measurement of endogenous REL2 protein in maternal sera samples. **(A)** Maternal sera (n=120) and control male (n=10) and female (n=30) serum samples were measured by IA-SRM with peptide qLYSALANK and two transitions (y6^1+^ and y7^2+^). A calibration curve was generated with the serial dilutions of recombinant mature REL2, light-to-heavy ratio for qLYSALANK and a sigmoidal curve fitting (**supplementary Fig. S6**). LOD (S/N>3) was determined at 9.4 pg/mL. As a result, endogenous REL2 in maternal sera was found at a median concentration of 331 pg/ml (IQR, 247– 444 pg/ml). A statistically significant difference between the three groups of samples was detected (Kruskal-Wallis test *P*=0.00001). **(B)** As measured by ELISA, endogenous REL2 was found at a median concentration of 1.6 ng/ml (IQR, 1.16 – 2.23 ng/ml). A statistically significant difference between the three groups of samples was detected (Kruskal-Wallis test *P*=0.000001). **(C, D)** Variability of median REL2 levels across gestational weeks, as measured by IA-SRM and ELISA, respectively. Error bars represent IQRs.

## DISCUSSION

Ten protein members of Insulin/IGF/Relaxin superfamily share structural similarities with insulin, including two short chains and a proteolytically-cleaved C-peptide^31^. Due to the distinct physiological roles as potent peptide hormones and high tissue specificity^1^, Insulin/IGF/Relaxin proteins warrant detailed investigation of their physiological roles and evaluation of disease biomarkers.

Due to immunoassay cross-reactivity and uncertainty about expression of low-abundance REL1_HUMAN and REL2_HUMAN proteins in human tissues and proximal fluids, the products of these two distinct human genes RLN1 and RLN2 were often collectively named a ‘human relaxin’ protein. Numerous studies previously investigated a ‘human relaxin protein’ and identified its role as a female and male reproductive hormone, a neuropeptide within the central nervous system, vasodilator, and cardiac stimulant in the cardiovascular system, and antifibrotic agent^32-34^. Expression of RLN1 gene was observed by RT-PCR in human prostate tissues and LNCaP prostate cancer cell line.^35-37^ According to the Human Protein Atlas, RLN1 transcript was classified as an exclusive prostate-specific transcript among the top 12 genes with highly enriched expression in prostate tissues, alongside KLK3, KLK4, and ACP3 genes. RLN1 was classified as an exclusive prostate-specific transcript, and the corresponding protein was suggested as an essential factor of male fertility^18,38^ and prostate growth^27^, and a prostate cancer biomarker^26,27,39^. A collective ‘human relaxin’ protein has been found in prostate and seminal fluid at concentrations of 8-50 ng/ml^17,18,26,40^. REL2 protein was predominantly expressed by the corpus luteum^31,41^, and was extensively studied as a pregnancy hormone^8^, with the levels exclusively measured by immunoassays. REL2 plays a crucial role in preparing the birth canal for parturition^31,42^, and is elevated in the first trimester^30,43^ reaching its peak concentrations of 2.2 ng/ml^30,41^. Similar to alpha-fetoprotein, levels of REL2 in the subsequent second trimester decreases to 1-1.3 ng/ml^30,44,45^. Low levels of REL2 during pregnancy have been associated with recurrent miscarriage^41^, toxemia of pregnancy, spontaneous abortion, ectopic pregnancy, and premature labor^44-46^. Those observations suggested that serum REL2 levels could serve as a potential marker of pregnancy complications, such as preeclampsia^47^.

RLN3 gene, another member of the relaxin family, is evolutionary distinct from RLN1 and RLN2, is predominantly expressed in the brain, and is involved in stress response, memory, and appetite regulation^48^. REL3 protein has been detected at levels ranging from 69 pg/mL^49^ to 35,000 pg/mL^48^. Such a broad range justifies the need for independent interference-free assays to quantify REL3, and the presented IA-SRM assays could offer a viable solution.

Proteomics by mass spectrometry is a promising tool to identify, quantify and characterize human proteins^50-52^. We previously demonstrated numerous applications of shotgun and targeted mass spectrometry to identify and quantify human proteins^53-63^, discover protein biomarkers^64-72^, and investigate the functional roles of human proteins^73-80^. The emerging proteomic assays combining the high analytical sensitivity of immunoassays with the nearly absolute analytical selectivity of mass spectrometry^81-83^ could resolve the issues related to the identification and quantification of structurally similar protein isoforms and proteins, including human relaxins. As independent assays, IA-SRM could detect and resolve the long-standing issues and cross-reactivities of ELISA. Analytical proteomics and the high-quality assay to measure low-abundance proteins in blood serum and proximal fluids are indispensable tools to verify and validate novel biomarkers, including presumably highly prostate tissue-specific genes such RLN1. The knowledge of the reference values and ranges of proteins in healthy population is imperative to enable identification and validation of novel disease biomarkers^84-86^.

In this study, we hypothesized that a highly prostate-specific protein REL1 (RLN1 gene) could emerge as a novel prostate cancer biomarker which has never been previously validated. IA-SRM and ELISA were chosen as independent assays, as we previously demonstrated for the highly prostate-specific proteins KLK4 and TGM4, and a prostate cancer-specific TMPRSS2-ERG fusion protein^3-5^. Several biological samples including cancer cell lines, seminal plasma, female and male blood serum, and maternal sera samples were measured for the expression of endogenous REL1. Despite the use of numerous techniques and approaches, REL1 has not been detected by IA-SRM in any biological samples, including a few samples highly positive with immunoassay measurements. Our IA-SRM assay detected REL2 in maternal sera (up to 1.3 ng/ml), but not REL1. Despite previous association of REL2 with cancer cell migration in breast cancer, we revealed the undetectable levels of REL1 and REL2 proteins in MDA-231 and MCF-7 breast cancer cell lines^15,16^. The best tryptic peptide of REL1 (<u>CC</u>LIG<u>C</u>TK) revealed a substantial post-translational modification (N-term pyro-cmC)^87^, relatively poor signal and poor LOD in IA-SRM assay (460 pg/mL). While the endogenous REL1 could be expressed at levels below LODs of ELISA and IA-SRM, our extensive exploration of literature and recent data in proteomic databases (Peptide Atlas^12^, MassIVE^88^) may suggest that RLN1 mRNA may not be translated into a human protein^89,90^. Interestingly, RLN1 gene is found only in apes and humans and has emerged due to duplication of a relaxin gene into two distinct genes RLN1 and RLN2. As we previously demonstrated for a testis-specific protein TEX101^50^, some highly tissue-specific genes could be nonessential. The relatively recent evolution of a highly prostate-specific gene RLN1 could explain the lack of its function^31^.

Since REL1 protein proved to be undetectable using IA-SRM, we diverted our efforts to REL2 protein, whose levels were reliably measured by ELISA and IA-SRM in some samples, including maternal sera. We hypothesized that establishing baseline reference levels of REL2 in maternal sera and control female and male sera would facilitate further investigation of REL2 as a marker of female reproductive system diseases, including ovarian cancer.

REL2 protein has previously been detected at 3 pg/mL^91^, 10 pg/mL^11^ and 46 pg/mL^49^ in healthy sera. Our ELISA measurements revealed 1,600 pg/mL median serum REL2 levels in the second trimester, in agreement with the reported concentrations^30,41,44,45^. ELISA also demonstrated high levels of REL2 (∼3 ng/mL) in some control female and male samples (**Figure 6**).

To quantify the endogenous mature REL2 by IA-SRM, we selected two proteotypic peptides DSWMEEVK and QLYSALANK. We discovered that the N-term glutamine of QLYSALANK peptide of the endogenous REL2 chain B was nearly completely modified (∼98.6%) to pyro-glutamic acid (pyro-Glu, qLYSALANK). Interestingly, pyro-Glu modification is infamous for human immunoglobulins^92,93^ and could arise due to *in vivo* modification by glutaminyl cyclase^94^ or spontaneous cyclisation during proteomic sample preparation. Peptide DSWMEEVIK revealed poor correlation with levels of QLYSALANK, qLYSALANK, DSSLLFEEFK, and QSEAADSSPSELK (**supplementary Fig. S8**), as well as ELISA, and was not considered for REL2 quantification. Peptide qLYSALANK exhibited relatively high correlation, and its measurements revealed the median serum REL2 levels of 331 pg/mL in maternal sera (**Figure 6**). Interestingly, nearly all control female and male sera, including those samples highly positive on ELISA, had REL2 levels below IA-SRM assay LOD of 9.4 pg/mL.

Relaxin has previously been suggested as a potential marker of ovarian cancer ^43^ and therapeutic target of ovarian cancer^95,96^ and cardiovascular diseases^24,34,97^. Patients with serous adenocarcinoma exhibited a mean relaxin level of 78.1 pg/mL^43^, and such levels could be verified with our IA-SRM assay. Our findings may establish a baseline for future investigation of REL2 levels in disease and provide a robust assay for diagnostic applications.

## CONCLUSIONS

We developed IA-SRM assays for the quantification of human REL1 and REL2 proteins in numerous biological samples, including blood serum. IA-SRM revealed that ELISA measurements could result in overestimated REL1 and REL2 levels which could be explained by cross-reactivity and non-specific binding. Despite high mRNA levels in LNCaP cells and previous literature reports, the endogenous REL1 protein was undetectable. REL2 protein was quantified by IA-SRM in blood serum, and a 21.8 pg/mL cut-off level unambiguously distinguished maternal sera (median 331 pg/mL) from the control female and male sera (median at assay LOD of 9.4 pg/mL). High selectivity and sensitivity of IA-SRM will facilitate quantification of endogenous REL2 in clinical samples, paving the way to its comprehensive evaluation as a disease marker.

## Supporting Information

Supplementary Figures S1 to S8 and supplementary Tables S1 to S11 present REL2 identification in maternal sera, correlation of IA-MS assay with ELISA, and evaluation of REL1 and REL2 expression in cell lines, heavy isotope labelled peptides internal standards, and patient samples.

## AUTHOR INFORMATION

### Notes

The authors declare no competing financial interest.

## ACKNOWLEDGMENT

We thank Jack Moore and Yuan Yuan for access and assistance with Q-Exactive instrument.

## ABBREVIATIONS

ELISA: Enzyme-linked immunosorbent assay
IA-MS: Immunoaffinity-mass spectrometry
IQR: interquartile range
LC: liquid chromatography
LOD: Limit of detection
ppm: parts per million
SRM: Selected reaction monitoring.

