## Supplementary Figures for "Identification and quantification of human relaxin proteins by immunoaffinity-mass spectrometry"

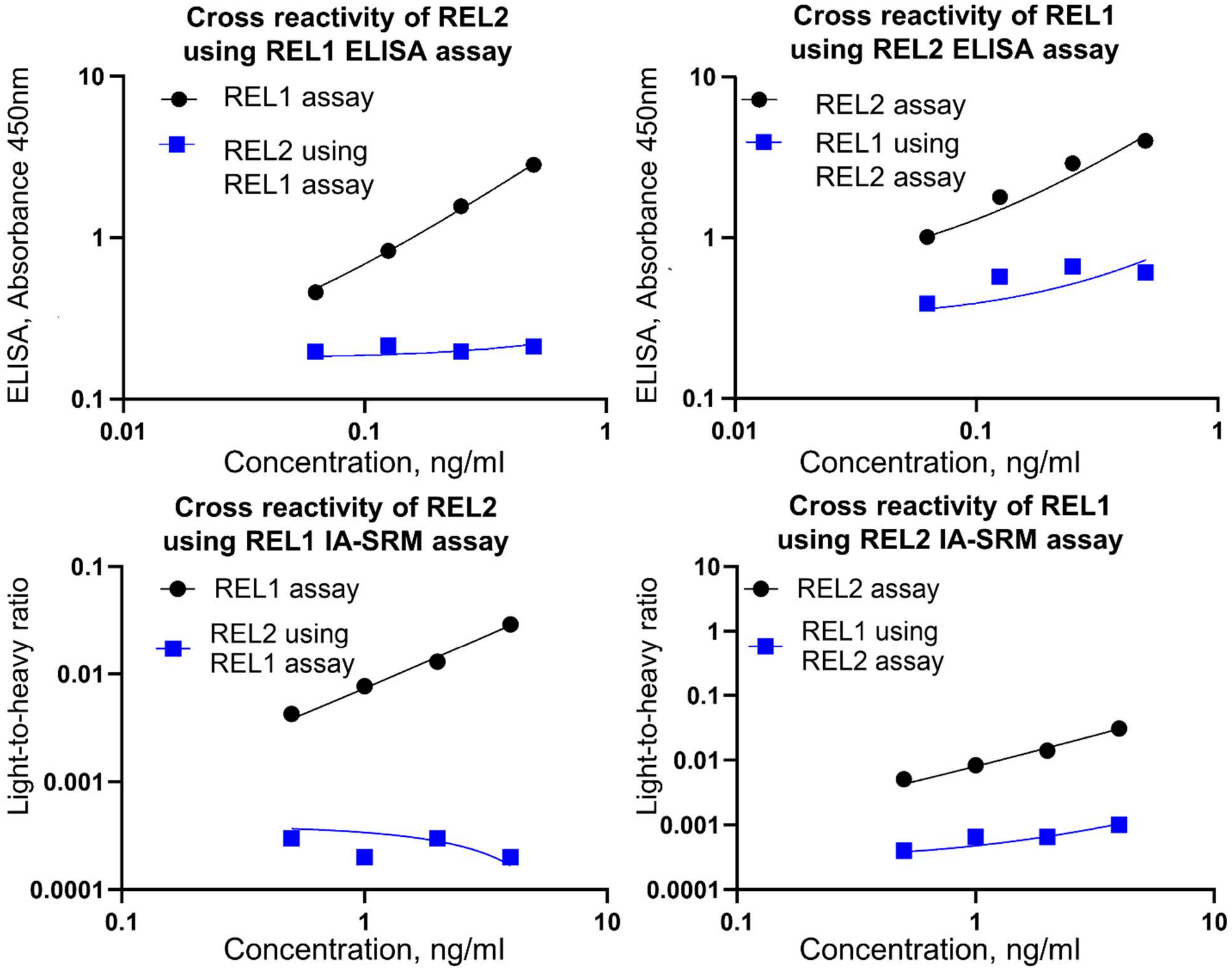


**Supplementary Figure S1. Cross-reactivity between REL1 and REL2 ELISA and IA-SRM assays.** To test for cross-reactivity, REL2 was used as the standard in the REL1 ELISA and IA-SRM assays. No cross-reactivity was observed. Similarly, no correlation was observed when measuring REL1 with REL2 ELISA or IA-SRM assay. The absence of cross-reactivity between the two ELISA assays and IA-SRM assays indicates the specificity of these assays in accurately distinguishing between REL1 and REL2.

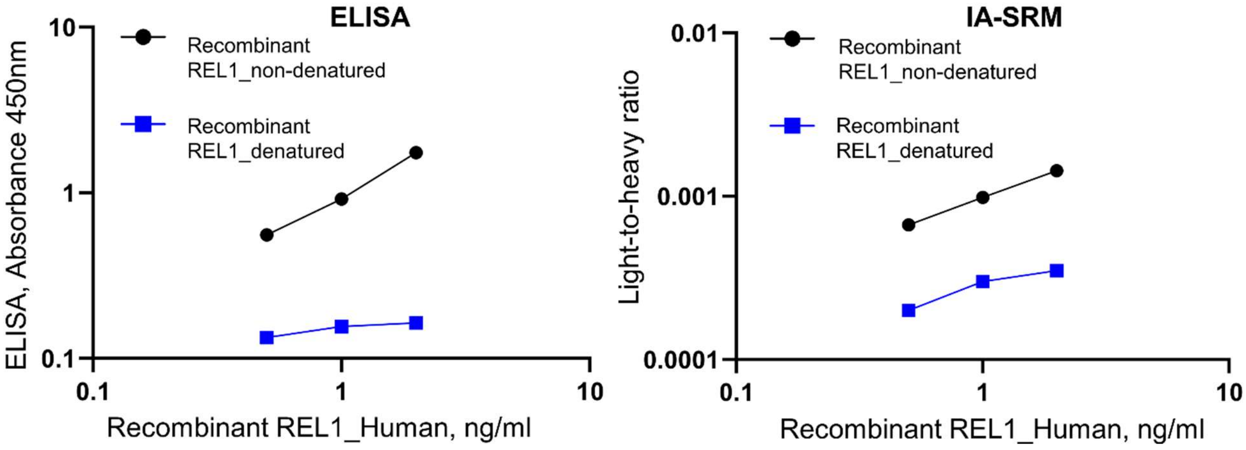


**Supplementary Figure S2. Recombinant REL1 measurement before and after denaturation using ELISA and IA-SRM assays**. Recombinant REL1 underwent denaturation through a process involving DTT, Rapigest, and heat treatment at 90 degrees Celsius. The complete loss of REL1 detection following denaturation indicates that the protein cannot withstand exposure to any such harsh treatments.

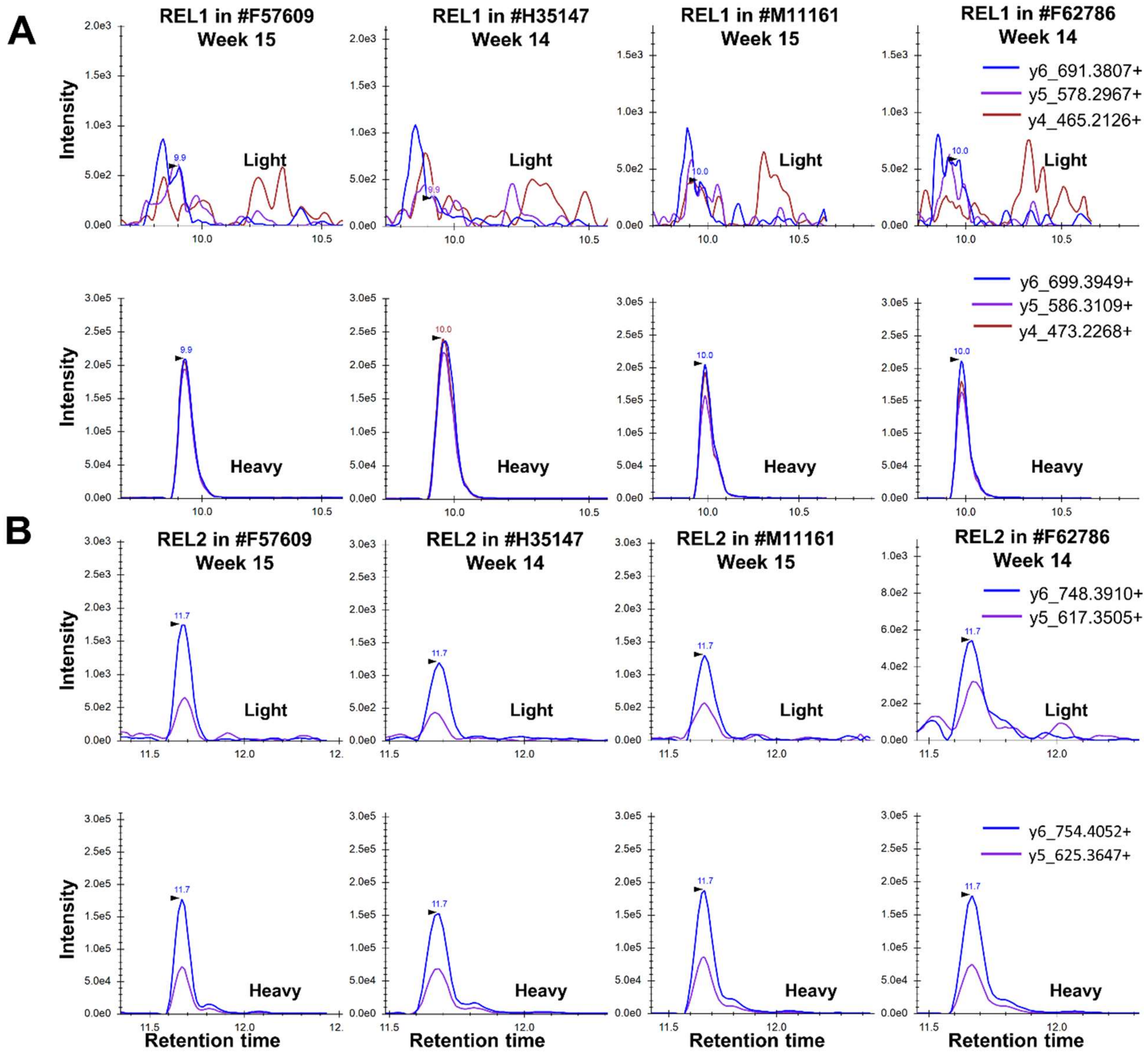


**Supplementary Figure S3. Measurement of REL1 and REL2 in four maternal sera samples using IA-SRM.** Initially four maternal sera samples, #F57609, #H35147, #M11161, and #F62786 were selected for the assessment of REL1 and REL2 expression. (A) REL1 was undetectable in any of the measured maternal sera samples. (B) REL2 was expressed in all four maternal sera samples.

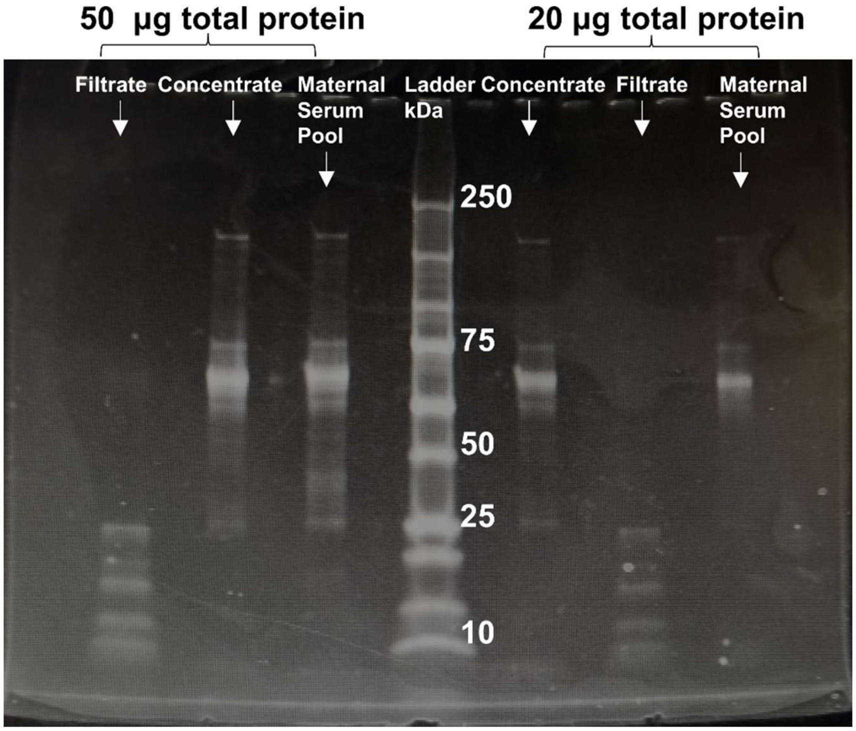


**Supplementary Figure S4. SDS gel illustrates the separation of high molecular weight (MW) and low MW proteins from the maternal serum pool using microcentrifuge ultrafiltration.** An amount of 50 µg and 20 µg of total protein of each of the filtrate, concentrate, and maternal serum pool were loaded onto the gel. Filtrate represents the solute that passes through the membrane. Concentrate represents the solute retained in the ultrafiltration tube. The maternal serum pool is constituted of 10 maternal sera samples expressing high levels of REL2.

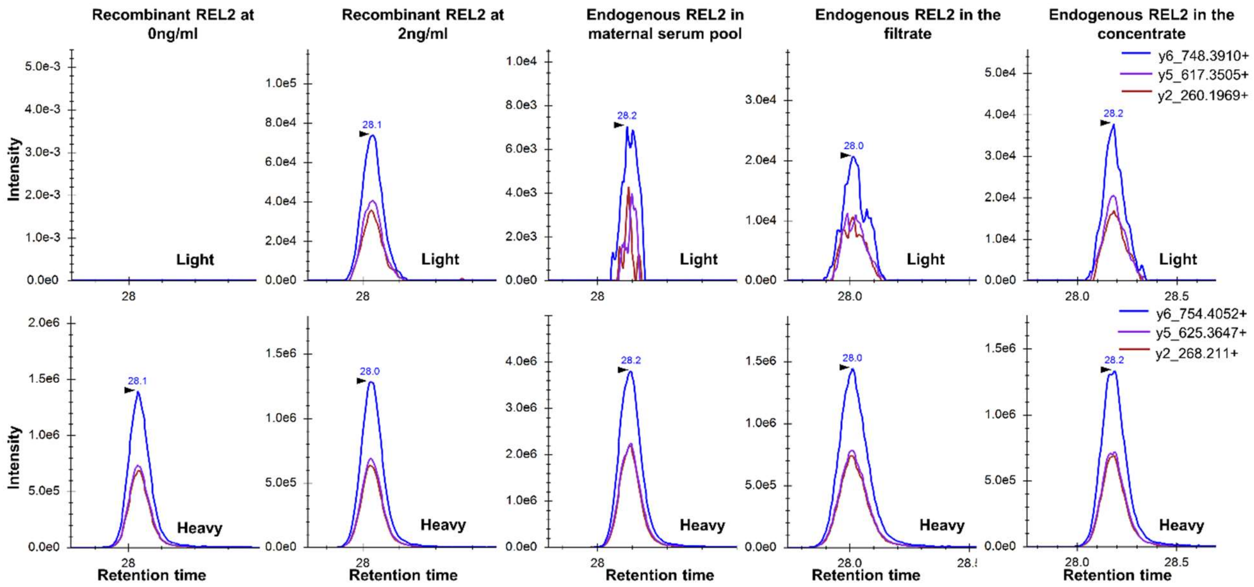


**Supplementary Figure S5. Preconcentration of endogenous REL2 by centrifugal ultrafiltration and lyophilization using SpeedVac.** Endogenous REL2 protein remained detectable in the maternal serum pool (before treatment), filtrate, and concentrate at different levels, suggesting 5-fold preconcentration. However, 5-fold preconcentrated REL2 was undetectable by shotgun orbitrap-ELITE, so no additional characterization of the endogenous REL2 (PTM modification, etc) was achieved. Recombinant REL2 at concentrations 0 ng/ml and 2ng/ml served as a negative and positive controls, respectively.

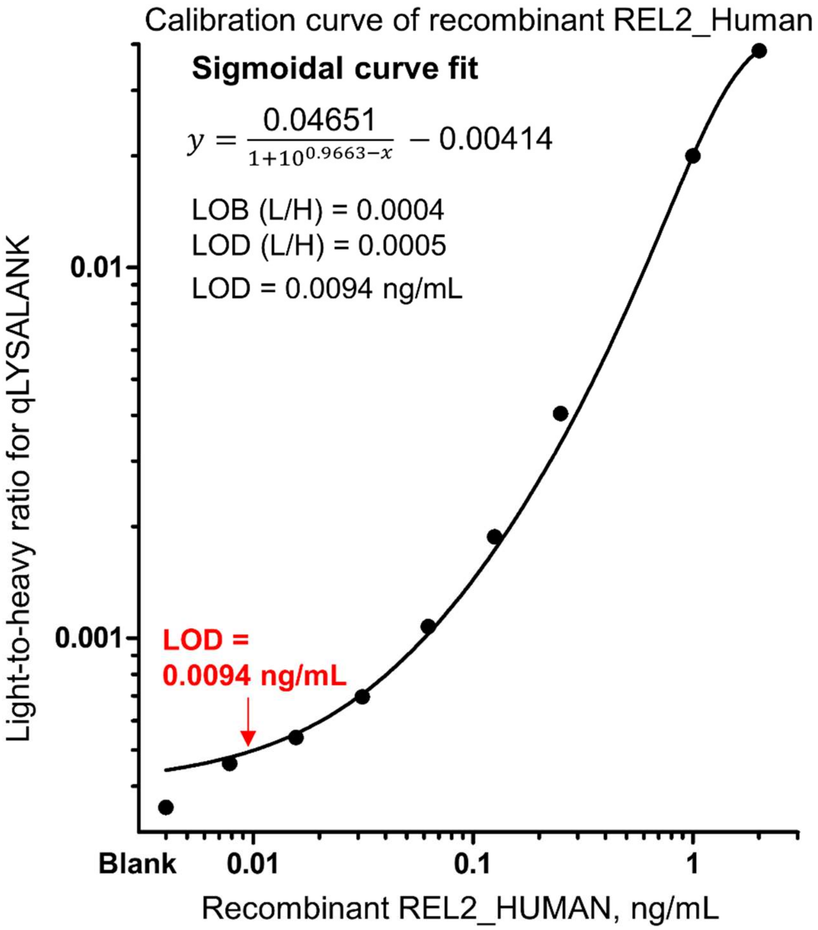


**Supplementary Figure S6. Calibration curve of recombinant REL2_HUMAN protein measured with IA-SRM assay.** Calibration curve was generated with serial dilutions of recombinant REL2 and measured with IA-SRM assay. LOD was determined at 9.4 pg/ml using light-to-heavy ratio for qLAYSALANK and sigmoidal curve fitting.

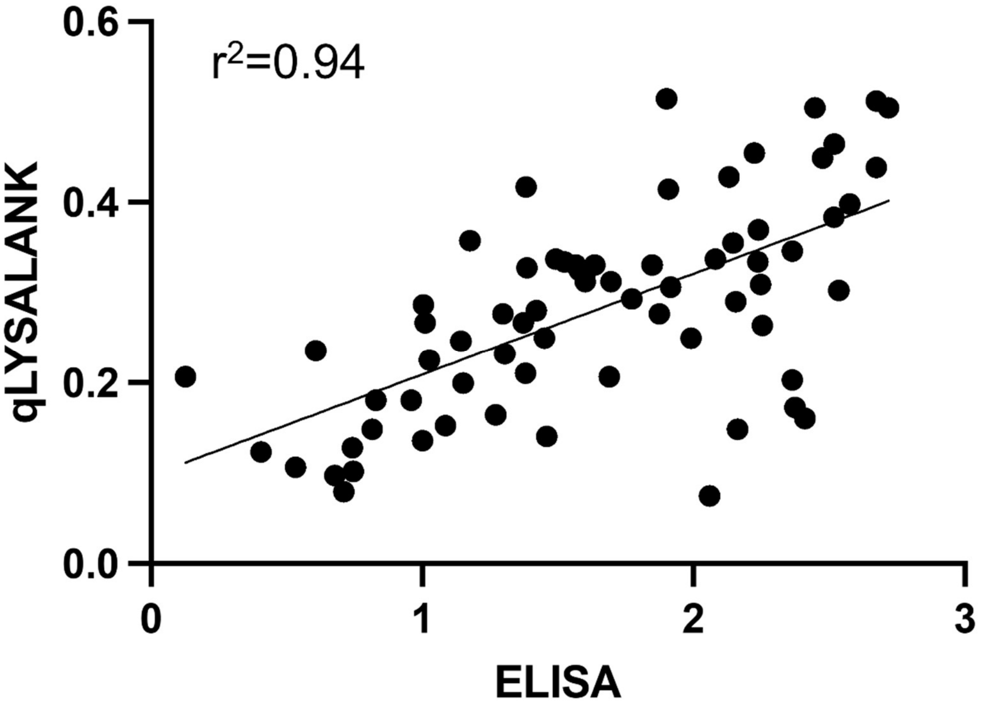


**Supplementary Figure S7. Correlation of endogenous REL2_Human protein measured in maternal sera samples using ELISA vs IA-SRM assay**. The tryptic peptide qLYSALANK and ELISA exhibited a high correlation, as indicated by $r^2$=0.94.

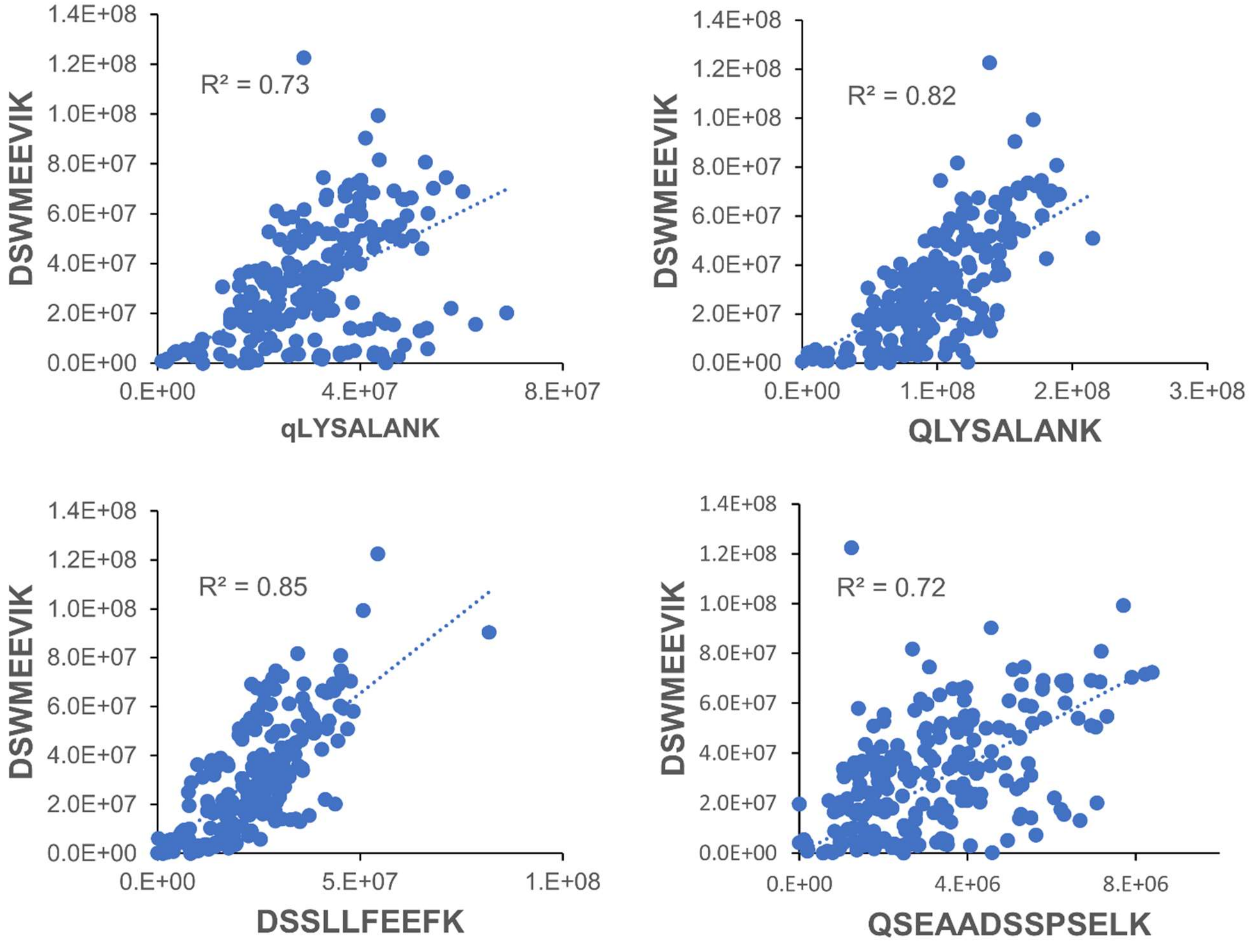


**Supplementary Figure S8. Correlation ($R^2$) of REL2 heavy DSWMEEVIK peptide to heavy qLYSALANK, QLYSALANK, DSSLLFEEFK and QSEAADSSPSELK.** Correlation of heavy DSWMEEVIK revealed poor correlation to all other REL2 tryptic peptides.
